# Just above chance: is it harder to decode information from human prefrontal cortex BOLD signals?

**DOI:** 10.1101/127324

**Authors:** Apoorva Bhandari, Christopher Gagne, David Badre

## Abstract

Understanding the nature and form of prefrontal cortex representations that support flexible behavior is an important open problem in cognitive neuroscience. In humans, multi-voxel pattern analysis (MVPA) of fMRI BOLD measurements has emerged as an important approach for studying neural representations. An implicit, untested assumption underlying many PFC MVPA studies is that the base rate of decoding information from PFC BOLD activity patterns is similar to that of other brain regions. Here we estimate these base rates from a meta-analysis of published MVPA studies and show that the PFC has a significantly lower base rate for decoding than visual sensory cortex. Our results have implications for the design and interpretation of MVPA studies of prefrontal cortex, and raise important questions about its functional organization.

## Introduction

The prefrontal cortex supports flexible, goal-directed behavior. Patients with frontal lobe lesions struggle to coherently organize their behavior around a goal and show reduced flexibility in changing circumstances ^1-3^. Theories of prefrontal cortex function emphasize its role in representing task-relevant variables like rules, goals, rewards, action choices, etc. during task performance ^4-10^. These task representations are hypothesized to serve as a source of top-down, contextual signals that bias processing in other brain regions, thus achieving cognitive control. Understanding the nature and form of prefrontal representations remains a key open problem in the study of cognitive control, learning, generalization, multi-tasking and decision making ^11-16^.

Much of our knowledge of prefrontal representations derives from single-neuron electrophysiology conducted in highly trained non-human primates. Such studies have consistently revealed rich coding of a variety of task-relevant information in the firing rate of individual prefrontal neurons ^17-21^, the activity patterns of ensembles of neurons ^22-26^ and in oscillatory synchronization of local field potentials ^27^.

In humans, blood-oxygenation-level dependent (BOLD) measures from functional MRI were traditionally seen as lacking the sensitivity and spatial resolution for the study of neural information coding ^28^. In the past decade, however, this view has been challenged by the development of sensitive multivariate pattern analysis (MVPA) methods that employ powerful machine learning pattern classifiers to decode the information content of spatially distributed BOLD activity patterns ^29-35^. MVPA has been applied to fMRI data from all over the brain, and the prefrontal cortex is no exception. Several studies have reported statistically reliable classification of task rule and other task-relevant variables from regions within prefrontal cortex ^36-41^.

An implicit assumption in many MVPA studies is that the function relating the information content of BOLD patterns with the information content of underlying neuronal activity is invariant across different parts of the brain. In other words, all regions have similar *base rates* for decoding information from BOLD patterns, to the degree that it is encoded in the underlying neuronal activity. For example, this assumption underlies analyses comparing decoding accuracies obtained from different brain regions, or those that employ roving, whole brain ‘searchlights’ to discover local regions that may carry such information. However, this remains an untested assumption. Indeed, the base rate for any brain region likely depends on interactions between the underlying micro-anatomy, neuro-vascular coupling and the raw signal-to-noise ratio, all of which may vary across regions.

For the prefrontal cortex, in particular, there is an impression among researchers with experience using MVPA that decoding information from BOLD patterns is particularly difficult. Despite the sensitivity of MVPA, typical group-mean classification accuracies reported in fMRI studies of PFC decoding often hover just above chance levels (e.g. values of 53%, 55% and 55% reported in Nelissen, et al. ^42^, Woolgar, et al. ^37^, and Bode and Haynes ^38^ for two-way classifications), even for task features like rules that are known to be robustly represented by the activity of prefrontal neurons in non-human primates ^20,22,23,43-46^. Consistently low classification accuracies hint at a low base rate for decoding information from prefrontal BOLD patterns. A low base rate may result, in part, from methodological factors, and in that case, it would be useful to know what these factors are. Alternatively, a low base rate may raise the possibility that the prefrontal BOLD signal itself may not adequately capture the information encoded in the spiking activity of prefrontal neurons. Such an observation would raise interesting theoretical questions about why and how prefrontal cortical coding differs in its type and organization from other parts of the brain with higher base rates.

A low base rate would also have implications for experimental design and inference. First, while a low base rate does not necessarily imply a small effect size (i.e. a low decoding accuracy may nevertheless be reliably different from chance), detecting a small difference would require that prefrontal MVPA studies be well powered. Second, a systematically lower base rate in PFC would complicate the interpretation of comparisons with other brain regions, which would require a consideration of the underlying base rates of each region.

In this paper, we empirically test the assumption that the base rate of decoding information from PFC is similar to other brain regions. To this end, we carried out a systematic meta-analysis of published fMRI studies of prefrontal cortex that employed MVPA. From this analysis, we estimate the base rate of decoding information from prefrontal cortex BOLD patterns and compare it to base rates obtained from visual cortex and mid-temporal regions. We also determine the distribution of classification accuracies obtained for ‘significant’ and ‘null’ effects in PFC and ask to what extent they overlap. Based on estimates of typical classification accuracies from these distributions, we also consider whether published studies typically collect sufficient data in order to detect small differences in decoding accuracy. Finally, we identify studies that have achieved considerably better-than-average classification accuracies and ask whether they are associated with particular sub-regions of prefrontal cortex, particular task features or particular analysis methods.

Collectively, our results show that the base rate of decoding information from PFC is just above chance levels, is systematically lower than other regions, and appears to be largely consistent across various methodological approaches. We conclude by considering the potential reasons for this low base rate of prefrontal classification.

## Results

### Typical decoding performance in prefrontal cortex is low

We leveraged our meta-analysis of published studies to approximately estimate the base rate of decoding information from PFC cortical BOLD patterns. To this end, we compiled all two-way, group-level mean classification accuracies reported across the 76 studies in our database. The resulting distribution is an estimate of the sampling distribution of mean classification accuracies for decoding information from PFC BOLD patterns. The mean of this distribution was 57.7% (95-CI: 56.3-59.2%), though we observed a skew, so that more than 63.1% of the accuracies were below the mean. Therefore, we employed the median as a measure of the central tendency, arriving at a base rate of 55.7% (95-CI: 55.0-57.0%). For comparison, we also derived base rates for decoding visual information from occipital and temporal cortex BOLD patterns. These were computed from meta-analytic data previously compiled by Coutanche and colleagues^47^. Compared to prefrontal cortex base rates, both the occipital and temporal cortex (median) base rates were significantly higher at 66.6% (95-CI: 61.5-72%) and 71.0% (95-CI: 68.0-75.0%) respectively.

MVPA studies of occipital and ventral temporal cortex focus exclusively on decoding information about visual stimulus attributes. This is because overwhelming evidence supports a strong prior for the hypothesis that the human occipital and ventral temporal cortices code for visual information. On the other hand, prefrontal MVPA analyses spanned attempts to decode a wide variety of information, reflecting a much less constrained hypothesis space for what information is represented in PFC. To control for this difference, we focused on a subset of 311 analyses in our database of prefrontal MVPA studies that attempted to decode “rule information”. Well-established deficits in rule-guided behavior have been linked to prefrontal dysfunction ^48-52^ and have been attributed to a loss of the ability to represent rules in working memory ^7^. Moreover, there is strong evidence from macaque electrophysiology that the prefrontal neurons code for task rules ^20,22,23,43-46,53^. Therefore, it is reasonable to place a strong prior on the hypothesis that task rule information is coded in the activity of human prefrontal neurons. A base rate obtained from rule decoding studies should, thus, be more comparable to the studies in Coutanche et al’s database. The median of the distribution of classification accuracies from rule decoding analyses was 57.5% (56.0-60.0%), again significantly lower than both the occipital and ventral temporal base rates.

These comparisons of base rates between visual and prefrontal cortex were made across different studies, likely employing different scanning methods, analysis pipelines, sample sizes, etc., which may all affect decoding accuracies. To address this, we examined prefrontal and visual cortex decoding accuracies obtained in a single fMRI dataset collected in our laboratory. fMRI data was collected while participants completed a cognitive control task that required encoding of both visual stimulus identity and visually-cued rules. This enabled us to compare decoding accuracies across regions while controlling for all other variables. We obtained a decoding accuracy of 55.4% for a two-way classification of rule in frontal cortex, and 72.3% for a two-way classification of stimulus-identity in visual cortex (Supplementary Figure S6, panel 4).

Finally, we leveraged our in-house dataset to determine whether differences signal-to-noise and pattern reliability mirrored these differences mirrored differences in decoding accuracies. While we found that raw SNR was actually higher in frontal cortex compared to visual cortex in our scanner (t=9.22, p<0.001), both functional SNR and pattern reliability were lower (t=3.99, p<0.001; t=11.91, p<0.001; Supplementary Figure S6, panels 1-3).

Collectively, these results demonstrate that the base rate for decoding information from human prefrontal cortex BOLD patterns is low in comparison to two sensory regions of the brain, consistent with the impression that MVPA in human prefrontal cortex is particularly difficult.

### Overlaps between decoding accuracy distributions for null and significant effects

Our database of PFC decoding analyses included group mean classification accuracies of both significant (greater than chance) and null effects. This allowed us to separately compile literature-derived distributions of classification accuracies for (a) when information is successfully decoded from frontal BOLD activity patterns and (b) when no information is detected. These distributions should overlap minimally if the studies that produced the classification accuracies had high power and low false positive rates.

The median of the classification accuracy distribution of significant effects was 58.7% (57.0-60.0%). Note that the values in our ‘significant’ distribution sample from an underlying ‘true’ distribution of decoding accuracies truncated at the left tail by the different significance thresholds used by specific studies. In other words, we are likely (conservatively) overestimating the center of this ‘true’ distribution. The distribution for the null effects had a median of 51.6% (CI: 51.0-52.0%). We computed the 95th percentile of this empirical ‘null distribution’ analogous to the typical ‘critical value’ used for null-hypothesis testing and obtained a value of 57.4% (CI: 56.0-62.0%). Note that these values sample a putative ‘true null’ distribution truncated at the right tail by the significant thresholds used in each analysis. Such truncating would normally bias our estimates of central tendency and the critical value downwards. On the other hand, the studies in our database employed different sample sizes and a variety of procedures for testing significance. If a proportion of these studies were underpowered, this would bias our estimates upwards. In addition, we have only relied on published studies in this meta-analysis. Thus, it is also likely that our estimates are biased upward due to the so-called *file drawer effect* or systematic non-reporting of null findings. In order to obtain a more conservative estimate of the critical value, we recentered the null distribution to 50%. With this approach, we obtained an estimate of 55.4 (CI: 53.5-60.1) for the 95th percentile.

Despite these conservative adjustments, as shown in Fig 2, there was considerable overlap between the estimated ‘significant’ and ‘null’ distributions. Indeed, 36.0% (CI: 15.5-66.4%) of the accuracies in the ‘significant’ distribution fell below the 95^th^ percentile of the uncentered null distribution of 57.5% while 23.5% fell below the centered null distribution of 55.4%. This overlap suggests that a number of previous studies were either not sufficiently powered to detect information coded in PFC BOLD patterns, or had inflated false positive rates (over the usual 5%).

**Figure 1.**
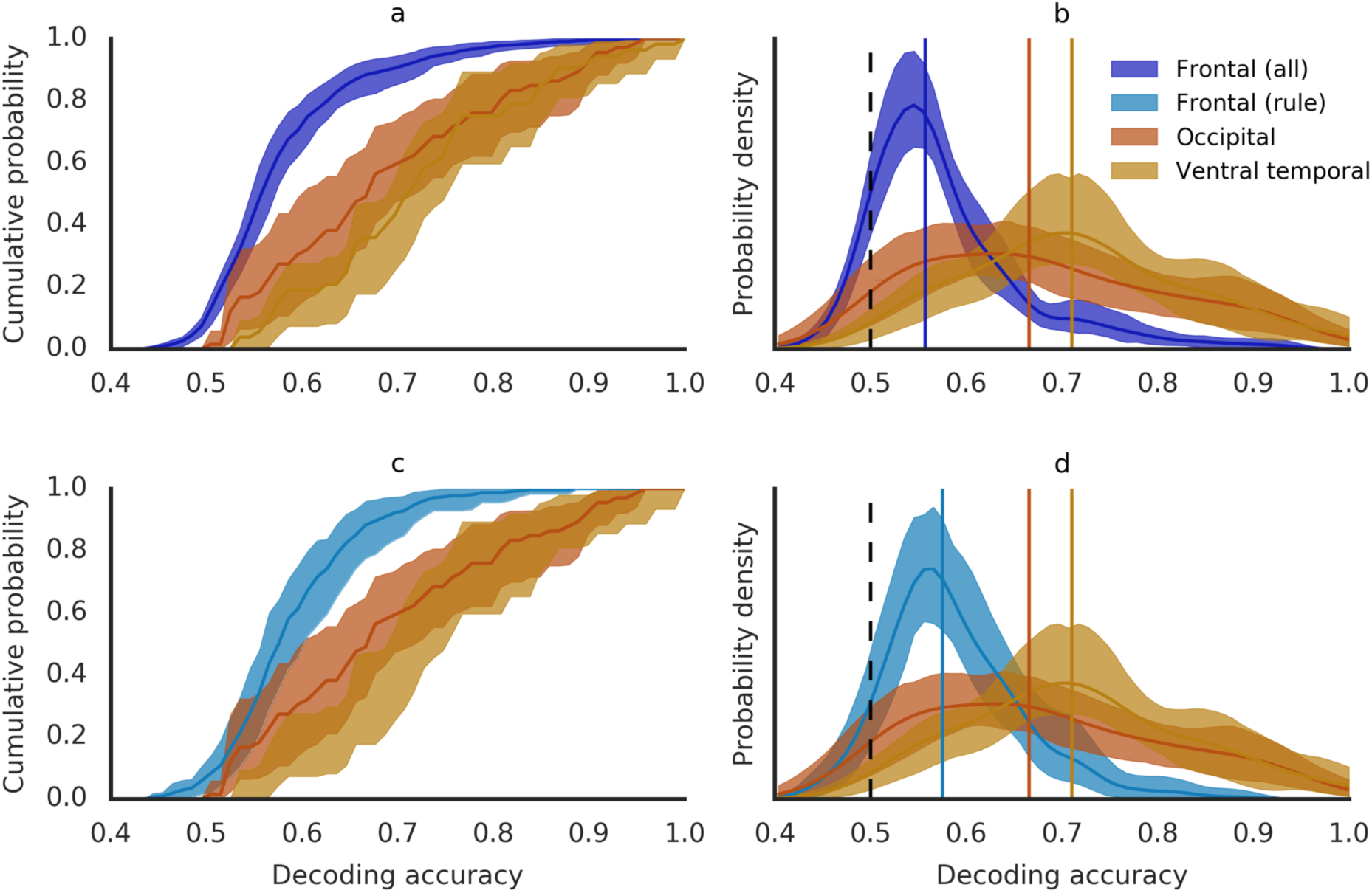
Decoding accuracy distributions for frontal, occipital and ventral temporal cortex. Cumulative distribution functions (a & c) and probability density functions (b & d) for visual decoding accuracies in occipital (red) and ventral temporal (orange) cortex compared with frontal decoding accuracies from all analyses (top panels, purple) and rule decoding analyses (bottom panels, aqua). Vertical lines indicate median values. Shaded areas reflect 95% confidence intervals obtained from a hierarchical bootstrapping procedure. Raw decoding accuracies are shown in Supplementary Figure S1, panels A & B.

**Figure 2.**
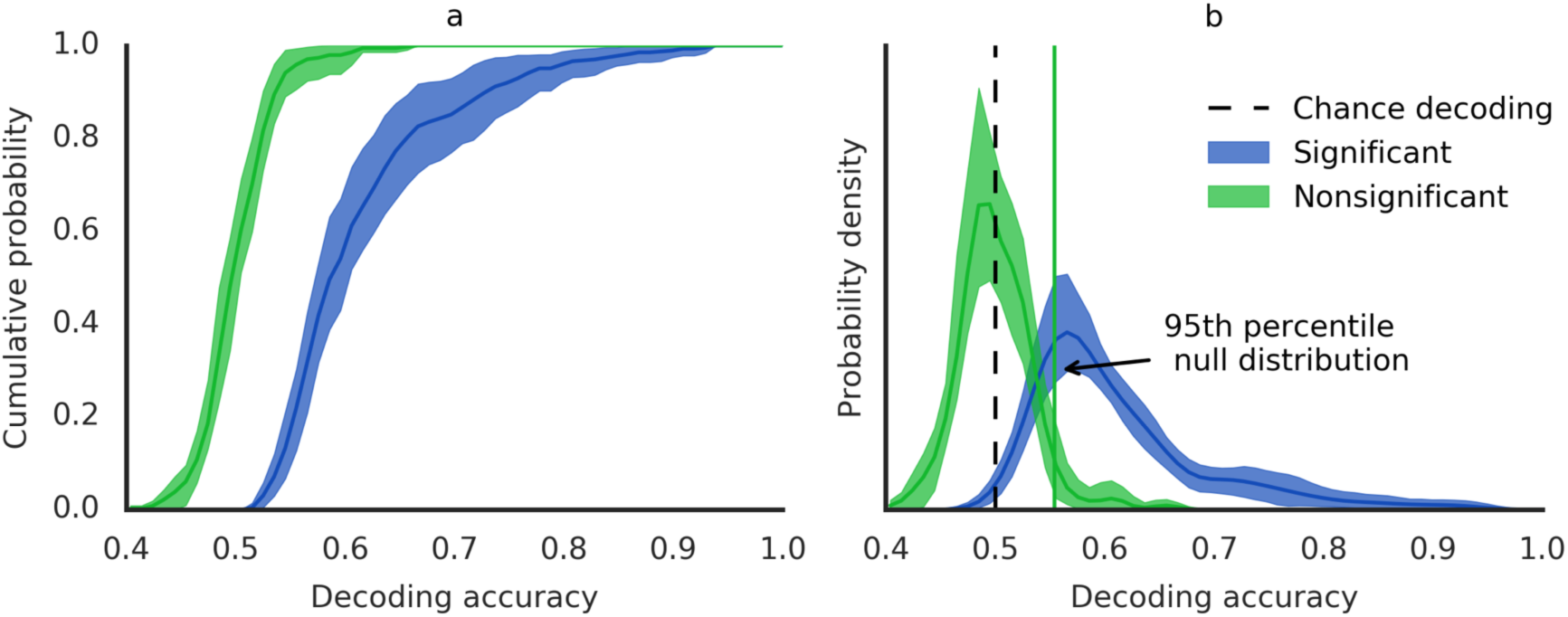
‘Significant’ v/s ‘Non-significant’ decoding accuracy distributions. Cumulative distribution function (a) and probability density function (b) for frontal decoding accuracies reported as significant (blue) and non-significant (green). Dotted line in (b) reflects chance-level (50%) and solid green line indicates 95^th^ percentile of the (centered) non-significant distribution. 23.5% of decoding accuracies in the significant distribution fell below this ‘critical’ value. Shaded areas reflect 95% confidence intervals obtained from a hierarchical bootstrapping procedure. Raw decoding accuracies are shown in Supplementary Figure S1, panel C.

However, as has been recently pointed out, the results of such tests of group-level mean accuracies against chance levels do not, in fact, support the population-level inference that the effect is typically present in the population ^54^. Instead, these tests assess the global null hypothesis ^55^ that *no participants* show the effect. As an example, in our lab’s fMRI dataset, the group-mean decoding accuracy of 55.4% was ‘significantly’ above chance in a parametric test against chance. However, only 5 out of 21 subjects showed a decoding accuracy that was significantly different from chance as determined by a non-parametric permutation test. Given this, it is critical to evaluate the prevalence of the effect. Indeed, it has been suggested that population inferences can be made based on the prevalence of an effect in a sample ^54^. As a consequence, the power to detect an effect at the level of an individual participant becomes particularly important.

It has not been common practice to report individual participant data, and therefore it is difficult to get estimates of within-subject variance needed for power analysis. Nevertheless, it is possible to make some general points about power at the individual level. The empirical ‘null’ distribution described above gives us an estimate of the typical decoding accuracy for a participant who does not show an effect (given that in these analyses the global null hypothesis cannot be rejected), and the 95^th^ percentile of the null distribution (55.4%) can serve as a rough estimate of a critical threshold which must be crossed for significance. On the other hand, an estimate of the typical individual’s decoding accuracy when they do show the effect may be derived from the ‘significant’ distribution. Comparing these two values will allow to estimate the magnitude of the difference in decoding accuracies that a typical study is trying to detect.

We consider two boundary conditions to derive the typical subject-level decoding accuracy. Consider the boundary condition where we assume that every significant effect reported in our database was maximally prevalent in that study’s sample (i.e. every participant shows the effect). In that condition, the median of the ‘significant’ distribution is a good estimate of the typical participant-level decoding accuracy. Given a median of 58.7%, we are, therefore, looking to detect a difference of only 3.3% points in order to reject the null hypothesis of chance-level coding for a typical participant. If a study included test 50 trials (which is typical for MVPA designs) for each condition, this would imply a difference of less than 4 trials successfully classified. Consider another boundary condition where every significant effect reported by studies in our database shows a prevalence of only 50% (given that a population inference requires that at least a majority of the participants show the effect). Assuming this liberal boundary condition for the studies in our database, we can estimate the typical decoding accuracy of an individual who showed the effect from the median of our significant distribution to be approximately 67.4% (assuming that half of the subjects in the study showed no effect and thus had a decoding accuracy of 50%). In that case, we would be looking to detect a 12% points difference – a difference of 12 trials successfully classified under the most liberal assumptions. These rough calculations suggest that the typical PFC MVPA effect is very small and given high levels of noise in fMRI measurements, one would require considerably more data per participant to detect such an effect reliably.

We emphasize again that we do not formally estimate effect sizes, which requires additional information of within-subject variance in decoding accuracies. Therefore, this analysis is not intended as a recommendation of trial numbers for future MVPA studies of the PFC and should not be cited as such. Nor do we recommend that the ‘critical’ value estimated above be generalized beyond this analysis to assess the significance of decoding accuracies in other studies. Rather, this analysis merely makes concrete the point that given the low base rate decoding accuracy in PFC and given the importance of assessing prevalence, sufficiently powered studies at the individual participant level are essential. Power calculations should be based on estimates of effect sizes from decoding of PFC BOLD patterns and not based on samples or effect sizes observed in other regions of the brain, given the differences between regions.

## Analysis of outliers

Our estimate of a literature-derived decoding accuracy distribution provides a means of placing the results of any given prefrontal decoding analysis within the context of the wider literature. Studies with very high accuracies, for instance, merit attention as they may have identified classes of information that are particularly well represented in prefrontal cortex, or may have employed a particularly effectively analysis approach. At the same time, given prior findings, these effects are surprising and, therefore, also merit closer scrutiny and replication to ensure that these results are not caused by other confounding factors. For these reasons, we examined the top 5% of reported classification accuracies in our database (Table 1) to identify factors that might explain the high values.

**Table 1:** Analyses in the top 5^th^ percentile of the ‘significant’ distribution

| Study ID | No. of analyses in top 5% (total 41) | Decoding accuracy range | ROI or Searchlight | Region(s)<sup>1</sup> | Description |
| --- | --- | --- | --- | --- | --- |
| 48 | 13 | 76% - 93% | ROI | Right precentral gyrus; Left precentral gyrus; Left middle frontal gyrus; Right supplementary motor area; Left supplementary motor area | Classified hand v/s saccade response |
| 66 | 5 | 79% - 90% | ROI | Right middle frontal gyrus; Left middle frontal gyrus; Bilateral anterior cingulate gyrus | Classified anticipation/experience of electric shock on arm v/s leg |
| 39 | 5 | 76% - 82% | Searchlight | Right superior frontal gyrus; Right anterior cingulate gyrus; Bilateral medial frontal gyrus | Classified to-be-purchases objects v/s neutral objects when attended/unattended. |
| 52 | 5 | 76% - 87% | Searchlight | Bilateral inferior frontal gyrus, pars orbitalis; Bilateral inferior frontal gyrus, pars triangularis; Bilateral inferior frontal gyrus, pars opercularis | Classified speech/music v/s re-ordered speech/music |
| 70 | 4 | 78% - 88% | ROI | Left middle frontal gyrus; Right middle frontal gyrus; Left precentral gyrus; Right precentral gyrus | Classified tasks that involved different rules & stimuli |
| 71 | 2 | 84% - 86% | ROI | Bilateral superior frontal &<sup>2</sup> gyrus & bilateral middle frontal gyrus & bilateral medial frontal gyrus & bilateral supplementary motor area;<br>Bilateral superior frontal gyrus & bilateral middle frontal gyrus & bilateral medial frontal gyrus | Classified reactive v/s predictive eye-movement pursuit of stimuli |
| 69 | 2 | 80% - 85% | ROI | Bilateral anterior cingulate gyrus & bilateral midcingulate area | Classified near v/s far semantic similarity of presented words |
| 54 | 2 | 81% - 83% | ROI | Bilateral precentral gyrus | Classified execution v/s imagining/observing reaching movements |
| 50 | 1 | 85% | Searchlight | Left inferior frontal gyrus, pars triangularis | Classified speech v/s spectrally-rotated speech |
| 31 | 1 | 77% | ROI | Left middle frontal gyrus | Classified prospective Yes v/s No decisions across intentions (honest/dishonest) |
| 78 | 1 | 79% | ROI | Bilateral precentral gyrus | Classified whole hand grasping v/s reaching movement |

1. Regions were registered to corresponding AAL region.
2. & refers to the union of AAL regions for constructing a larger ROI used in the corresponding study.

First, as many as 18 of the 41 analyses in the top 5% decoded some form of motor response (reaching, grasping, saccades etc.). Five additional analyses involved classifying the anticipation or experience of electric shocks on very different parts of the body (arm v/s leg). Six other analyses classified ordered stimuli like speech or music versus unordered versions of the same stimuli. All of these studies manipulate conditions that likely produce univariate differences, either as a small mean-response difference across a majority of the voxels or differential activation of adjacent subregions. For example, univariate analyses of ordered vs unordered stimuli contrasts are often used to localize language specific regions in prefrontal cortex ^56,57^. Univariate contributions to decoding analyses do not invalidate the inference of information coding. But, they do not require the use of pattern classifiers to detect, and therefore, should not inform an assessment of the method as it is used in more typical MVPA analyses.

Two studies employed unusual measurement or analysis methods. Study 39 (5 analyses) deployed non-linear classifiers which produced significantly higher accuracies than linear classifiers on the same data. The classification accuracies obtained from the linear classifiers are much closer to the median of our distribution. Study 31 uniquely employed high-resolution scanning with a 7T magnet.

Importantly, only 4 of these analyses, all from a single paper (Study 70), involved decoding task or rule information as posited by models of cognitive control and observed in macaque studies. However, even in these analyses, task/rule was confounded with visual information as the contrasted task conditions involved different classes of stimuli. Indeed, when task/rule was decoded after controlling stimulus differences, classification accuracies were in the mid 50s, close to the median of our distribution. Therefore, the high classification accuracies may have been driven by the additive effects of multiple sources of information.

In summary, we did not find a particular factor or approach that consistently explained these outlying decoding accuracies beyond what basic univariate analysis could provide.

### Factors affecting decoding performance

We next sought to systematically examine whether particular sub-regions of frontal cortex, particular types of information, or particular methods were correlated with decoding accuracy levels. To do this, we assessed the partial influence of these factors in our full database using mixed-effects linear regression. We fit a single mixed-effects regression model with all the characteristics - region, information type, analysis procedure. All these regressors were dummy coded with one category omitted from the model. To account for covariance between observations from the same study, we also included random study intercepts. Classification accuracies reported as non-significant were excluded as they are more likely to have small or no effects.

To assess the significance of each characteristic, regressors were dropped one at a time from the model and tested against the full model using the likelihood ratio test. Results are shown in Fig. S2. Only the inclusion of *classifier* significantly improved model fit (Supplementary Table S4; *L*=17.3 *p*=0.004). Post-hoc pair-wise tests applied to *classifier* showed that non-linear SVM had a significantly higher accuracy than linear SVM (*p*=0.02 Tukey-HSD). This effect was driven by 3 out of 4 studies using non-linear SVM, each with accuracies above 70%, and was also significant in a regression using only a single mean accuracy per study (Supplementary Table S5, *F*=4.14, *p*=0.003). One of these studies were also identified by our outlier analysis. However, given the non-normal distribution of group accuracies, this result should be evaluated with caution. The effects on accuracy for all other analysis characteristics can be seen in Supplemental Figure S2 and Supplemental Table S4.

The full regression model did not reveal differences in accuracy across regions. Our regression analyses may have been underpowered due to the large number of regions tested simultaneously and the few number of observations associated with each region. Given that we found no effect of ROI laterality in the main model, we combined the left, right and bi-lateral portions of each ROI for a more powerful, exploratory follow up analysis, and observed that superior & middle frontal gyrus, orbital part (58.2%), and middle frontal gyrus (59.7%) were marginally lower than the grand mean (61.3%) (Supplemental Figure 3; *t*=-2.01, p=0.044; *t*=-2.21, *p*=0.027). Next, we tested whether accuracy differed across regions based on the type of information decoded. Not surprisingly *response* decoding was associated with higher accuracy in superior frontal gyrus (66%, *p*=0.03) and marginally so in precentral gyrus & supplementary motor area (64.3%, *p*=0.09), and *perceptual* decoding was associated with higher accuracy in cingulate cortex (68.5%, *p*=0.004; Supplemental Figure S4). There were no other differences in accuracy across the rest of the regions. Finally, we carried out a second regression that focused only on frontal regions within the multiple-demand (MD) network^58^ that response robustly to manipulations of various cognitive demands. These ROIs are more representative of typical functional clusters observed in fMRI studies of frontal cortex (as opposed to the large AAL ROIs), and may better reflect frontal decoding properties. However, we again found no differences within MD regions, or between MD and other regions (Supplementary Figure S5). In summary, although these different analyses of our data suggest small differences across regions - OFC, motor, cingulate – exist, the distribution of classification accuracies is broadly similar and low across prefrontal cortex.

## Discussion

Over the past decade, MVPA has emerged as a powerful method for studying information coding in the human brain with fMRI ^29-34^. MVPA is a more sensitive method than traditional univariate analyses because it combines evidence across voxels to detect subtly encoded information in distributed patterns of activity. Given the many open and important questions regarding nature and form of prefrontal cortex representations, MVPA has also been enthusiastically applied to this brain region. Previous studies have implicitly assumed that the base rate of decoding information from PFC BOLD patterns is similar to that of other brain regions, or at the very least, does not vary systematically. This assumption underlies the comparison of decoding accuracies from PFC with other regions of the brain, the use of whole-brain searchlights to discover regions of coding, as well as choices about sample size, analysis methods, etc. This assumption has not been tested before and contrasts with a prevailing impression amongst practitioners (often heard at conferences) that decoding information from PFC BOLD patterns is uniquely challenging. We estimated the base rate decoding accuracy for PFC and show that it is indeed lower than sensory (visual) cortex.

Our meta-analysis of prefrontal MVPA studies identified over 800 MVPA decoding analyses across 76 studies, each reporting a group-level mean classification accuracy. This dataset includes attempts to decode a wide range of information from BOLD patterns in various subregions of prefrontal cortex, while employing a similarly wide range of MVPA methods. In this sense, our meta-analysis samples a space of possible approaches to classification in PFC and so permits not only an estimate of a base rate, but also determine which particular approaches are systematically more or less successful.

From this dataset, we estimate the base rate for decoding information from prefrontal BOLD patterns at the low value of 55.7% for two-way classifications where chance performance is 50%. Further, we observed that the PFC base rate is markedly lower than base rates for decoding visual stimulus information occipital and ventral temporal cortex BOLD patterns which were at 66.6% and 71.0% respectively. These differences are not due to the larger hypothesis space of PFC decoding studies. The differences remain stark even when we derive the PFC base rate estimates solely from analyses that decode rule or task information, which we believe is very likely to be coded by PFC neurons given evidence from primate electrophysiology and human neuropsychology. Indeed, we may have underestimated the PFC base rate given that we only included studies that mentioned prefrontal cortex in the abstract. Other studies that ran whole brain searchlights and found chance-level coding in PFC may have been excluded as a result. In practical terms, this low base rate means that it is likely that the difference between studies reporting successful versus unsuccessful classification may hinge on only a few trials classified better than chance. This has significant implications for experimental design and inference that we discuss further below.

Further, this base rate provides an empirically derived prior against which future decoding paradigms or methods can be compared. For example, studies that propose a new feature of the fMRI signal (for e.g. Waskom and Wagner ^41^ recently decoded context information from local connectivity measures), or a new decoding method as capturing a special aspect of coding in the PFC, can be evaluated against this base rate prior. In other words, we can ask whether incremental gains in decoding accuracy, beyond those expected given the base rate, are achieved from applying the new feature or method. We emphasize, however, that any such comparisons of decoding accuracies must take into account the underlying variance in decoding accuracies (as estimated by appropriate non-parametric methods).

Similarly, our base rate provides a principled basis on which to highlight past studies that were unusually successful at decoding information for further scrutiny. We probed several outlier analyses in our data set that showed large classification accuracies to look for a consistent feature that explained their success. Most of these cases could be attributed to the influence of univariate effects. Beyond these effects, the few remaining outlier studies did not share a consistent approach or classification type that resulted in a marked shift in criterion. Nevertheless, our analysis places the likelihood of these outcomes in context given the broader literature. As such, these individual studies might merit further follow up and replication.

Beyond consideration of the outliers, we leveraged the meta-analysis dataset to ask whether particular information-types or methodological choices were consistently associated with higher decoding performance using regression. We found some evidence that motor information in some regions of posterior prefrontal cortex and perceptual information in cingulate cortex are associated with slightly higher decoding performance. Conversely, regions of mid-lateral PFC closely tied to cognitive control were associated with, if anything, even lower classification success than other areas of the frontal lobe. We also found a benefit of using nonlinear classifiers in the small number of studies that use them. However, this benefit may be offset by known complications associated with the use of non-linear classifiers. While they are indeed able to be able to read-out a wider variety of representational formats, non-linear classifiers are more susceptible to over-fitting. Moreover, a ‘linear readout’ (that a linear classifier implements) is often considered a hallmark of an *explicit* representation ^59,60^ under the assumption that downstream neurons usually implement a linear readout. Therefore, the results of a linear classifier can support stronger claims about representations that those of a non-linear classifier cannot. Nonetheless, the non-linear SVM approach may merit further study and replication of its advantage for PFC classification. Beyond this, we found that decoding performance was robust to variations in methods, with the one caveat that our power to detect these effects was not high. Collectively, these results suggest that the low base rate of decoding information from PFC BOLD patterns is a very general finding.

What makes decoding information from PFC BOLD patterns so difficult? Electrophysiology studies in the non-human primate have provided consistent evidence for ubiquitous coding of task-relevant information in prefrontal firing rates ^17-21^. Indeed, recent evidence suggests that macaque PFC representations of task variables are high-dimensional, and that this property enables these task variables and their conjunctions to be read out by a linear classifier.^24^ Thus, the finding of a low base rate of decoding such information from prefrontal BOLD patterns is surprising. Furthermore, the differences in base rates between prefrontal and occipital/ventral-temporal cortex suggest that the function relating the information content of spiking activity and that of BOLD patterns across voxels varies across regions. Of course, it is conceivable that human PFC representations have different properties than those of macaques and that prior research has simply not identified the appropriate contrasts or type of classifier to probe them. However, we deem this unlikely as a general account and suggest that PFC BOLD patterns across voxels may only weakly reflect the information encoded in the firing rate of populations of prefrontal neurons.

The base rate of decoding information is no doubt influenced by the properties of PFC representations. For example, an oft-cited feature of PFC neurons is that they display ‘mixed selectivity’^24^ or ‘adaptive coding’^6^ – i.e. their selectivity for particular task-variables is highly context and task dependent. Such a feature of coding may render population activity patterns more susceptible to noise contributed by uncontrolled features of the environment like temporal context, thus making decoding more difficult. Similarly, the activity of PFC neurons is known to show a greater degree of temporal autocorrelation^61^, which may heighten the similarity between condition-specific activity patterns. Finally, population representations of the contents of working memory in PFC are known to be highly dynamic^23^ with each to-be-remembered item producing a complex trajectory through neural state space, suggesting that information may be stored in the temporal profile of these trajectories, rather than only in overall activity. All of these properties likely affect the decoding of information, though note that they would affect decoding from electrophysiologically measured firing rate patterns as well, not just BOLD patterns. Therefore, they are not sufficient to explain the particular difficulty with decoding from BOLD patterns.

Low decoding based rates in PFC may be caused by MR-induced or physiological noise contributions to the BOLD signal that may influence the trial-by-trial variability of BOLD patterns in a region-specific manner. In our own fMRI dataset, empirical estimates of raw signal-to-noise ratios were not lower in PFC compared to visual cortex. Therefore, the lower decoding accuracies we found in PFC could not have been driven by raw noise differences. However, both, the univariate functional SNR, and the reliability of BOLD patterns were lower in prefrontal cortex than in visual cortex (Supplementary Figure S6). A lower reliability of BOLD patterns in PFC would certainly make decoding more difficult. However, this lower reliability also demands explanation, and would be affected by the other factors we discuss.

Another possibility we find more likely is that the particular, local functional organization and distribution of neural populations in prefrontal cortex may reduce differences between conditions at the voxel scale measured with fMRI.^62^ In a recent study, Dubois et al. (2015) examined the coding of face viewpoint and identity information using both MVPA of BOLD patterns and single-unit recordings in macaques. While both viewpoint and identity were strongly coded in single-unit firing rates, MVPA of BOLD patterns only revealed viewpoint information. The authors concluded that identity decoding suffered because identity coding neurons were only weakly clustered spatially as compared to viewpoint-coding neurons. Clustering may enable nearby blood vessels to be strongly driven by neurons selective to one condition, thus enabling inhomogeneities in the sampling of the activity of selective neurons by voxels ^63^. Most single-unit studies in primate prefrontal cortex, however, show very little evidence of clustering ^64^, with neurons coding different task-relevant information being heterogeneously intermixed at a fine scale e.g. ^21,65,66-69^ By contrast, in the visual cortex MVPA effects may depend on clustering both at the fine-scale in the form of columnar structure ^63^, and also at the coarse-scale in the form of asymmetric spatial distribution of columns ^70-72^. This interpretation suggests that higher resolution fMRI might ultimately help this base rate issue, though this might require still higher resolution than is currently feasible. Alternately, PFC representations may perhaps be better studied by leveraging repetition suppression effects ^73,74^, which are not affected by the local distribution of neural populations.

Regardless of the source of these differences, a base rate decoding difference between prefrontal and visual cortex has important implications for the interpretation of studies which rely on comparisons of classification accuracies across regions. Consider, for example, the debate surrounding the locus (prefrontal or sensory cortex) of detailed sensory information during working memory delays, which has been informed by the finding that, while classifiers readily decode sensory information from the BOLD signal recorded from visual cortex, they are much less successful in the frontal cortex reviewed in ^75^. If the base rate for decoding is lower in prefrontal cortex, such a finding, on its own, would provide limited support for an exclusive sensory cortex locus of working memory representations. In order for such comparisons to be interpreted, it would be critical to first consider the base rate for decoding information for the regions in question. Indeed, while we have focused on the PFC in this study, our results also highlight the more general point that knowing such base rates is critical to interpreting the findings of any MVPA study, including those employing other measurement modalities that span the brain like MEG or EEG

The empirical distribution of PFC classification accuracies that we have compiled allows any new result to be placed within the context of prior findings and for its likelihood to be computed. As also noted above, analyses that report unusually high classification accuracies should draw attention for the possibility that they may be false positives or be driven by confounding factors. Employing this logic, we compiled separate distributions of classification accuracies for significant and null effects from our dataset and observed considerable overlap between these empirical distributions. This overlap suggests the presence of a number of analyses that either did not appropriately control false positive rates, or were insufficiently powered to reject null hypothesis of chance-level coding. Indeed, we noted the widespread use of parametric statistics, which have been shown to inflate false positive rates for classification accuracies. We agree with recommendations that MVPA studies should rely primarily on appropriately conducted permutation testing at the individual level ^76^ and the assessment of the prevalence of effects at the group level ^54^.

Indeed, given the importance of the prevalence of MVPA effects in making population-level inferences ^54^, it is important to consider the power of an experimental design to detect an effect at the individual-subject level. Based on our rough estimates of typical decoding accuracies from the ‘null’ and ‘significant’ distributions, we expect that significantly more data *per participant* will need to be collected to detect small differences in decoding accuracies more consistently. This is particularly important as future studies move beyond demonstrating information coding to examining the factors that may influence the properties of underlying representations. With classification accuracies typically hovering in the 50%-60% range, there is little room to detect their modulation with experimental manipulation or by incorporating covariates without many more measurements. Similarly, improved statistical power will also be necessary in order to regress out the potentially confounding effects of small, idiosyncratic differences between task conditions on nuisance variables like difficulty or time-on-task ^77^. The prefrontal BOLD signal is known to be sensitive to such variables ^56,78^ and regressing out their effects post-hoc is critical to unbiased inference.

In conclusion, we provide an estimate of the base rate of decoding information from PFC BOLD patterns and show that it is markedly lower than two brain regions in visual cortex. Our low estimate supports the prevailing impression that using MVPA to decode information in PFC is particularly challenging. The reasons for this difficulty remain open, and we suspect may reflect an important property of neural coding in the PFC, such as their spatial organization and distribution. Though we cannot pinpoint the specific factor driving this difference, our results have concrete implications for the design and interpretation of future studies – we recommend more data per participant, the use of permutation tests, reporting of prevalence at the group-level, and a consideration of base rate when making comparisons across regions. Finally, this study provides an example of how meta-analyses of MVPA data can provide unique insights that are not available in single studies. To facilitate future investigations, we are sharing our database and code publicly via the Open Science Framework (OSF) (https://osf.io/8dvzr/).

## Methods

### Literature Search and Study Inclusion

We conducted a comprehensive search of the literature to identify all published studies between the years of 2001 and 2016 that employed multivariate methods to decode information from fMRI BOLD patterns in the prefrontal cortex. In summary, we queried the PubMed database for articles whose abstracts contained at least one term related to (i1) functional imaging, (i2) multivoxel pattern analysis, (i3) frontal cortex. In addition, we explicitly excluded articles whose abstracts contained terms related to patient samples and non-human primates. A full list of terms employed for the search are shown in Fig 3.

**Figure 3.**
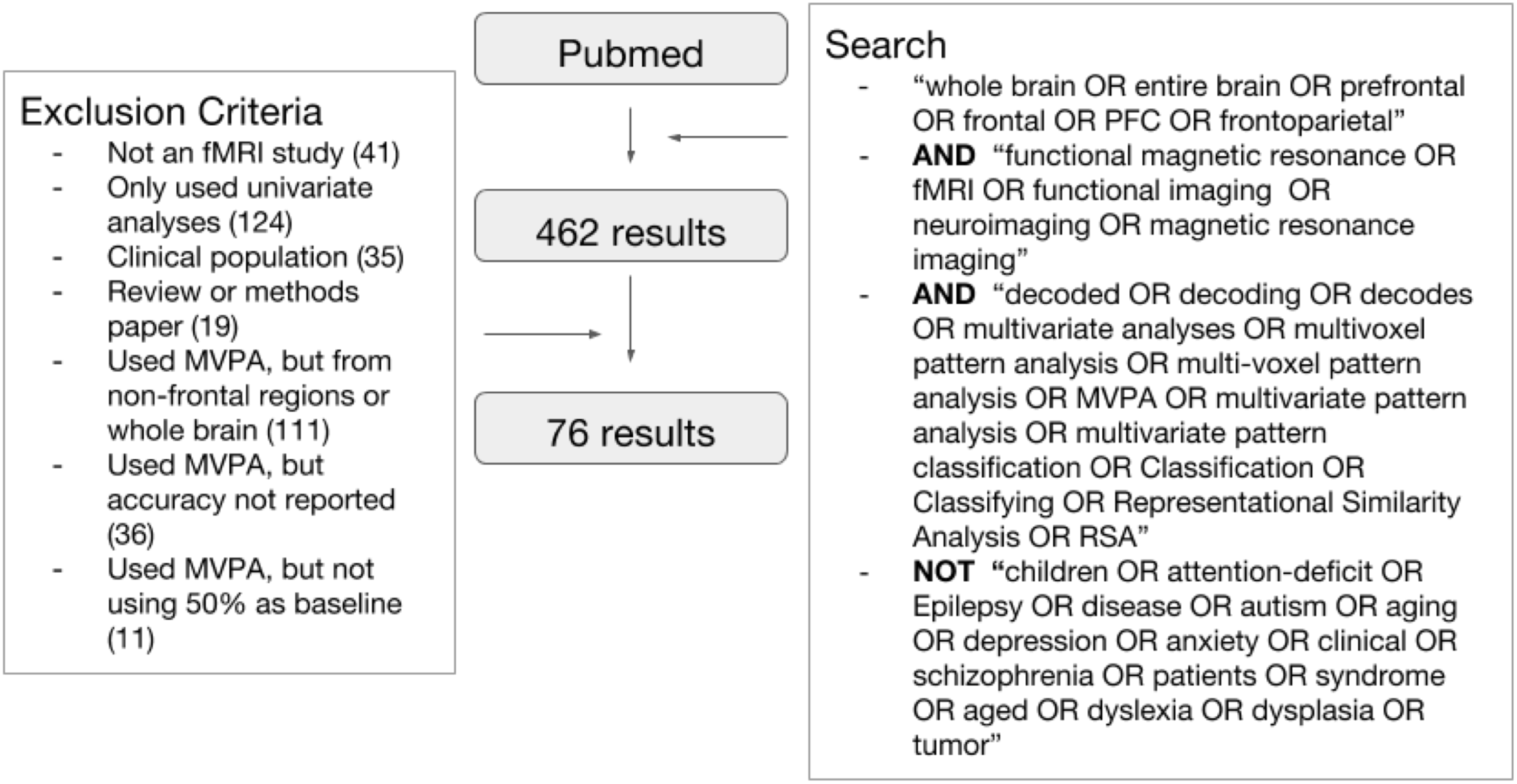
Search terms employed for literature search. The final literature search was conducted on 09/03/2016. The search string above was entered into Pubmed’s advanced search (https://www.ncbi.nlm.nih.gov/pubmed/advanced), additionally restricting the year of publication to be between 2001 and 2016.

This search resulted in a set of 462 studies that employed multivariate fMRI analysis, including classification analysis and representational similarity analysis. Across approaches, there was further variability in the metrics used to report the strength of decoding, including significance statistics, correlation coefficients, single-subject mean classification accuracies, group-level mean classification accuracies, etc. To allow for comparison and aggregation across studies, we focused on the largest subset of studies, those that employed a cross-validated classification approach for decoding and reported group-level summary classification accuracies. We define the cross-validated classification approach as one in which unseen and unlabeled BOLD patterns are assigned labels by a ‘classifier’ trained on independent data, and the success of this classification is reported in terms of classification accuracies. We further restricted our dataset to the studies that reported decoding analyses with two classes (i.e. those that had 50% as chance level). This left us with 76 studies, the list of which can be found in Supplementary Table S1.

### Within Study Extraction

Studies reported a variable number of decoding analyses, ranging from 1-96 per study Some studies decoded the same information from different PFC regions, while other studies decoded different types of information from the same region. A few studies also applied the same analysis to data from different time points within a single trial. As a rule, we separately recorded all reported group-level summary classification accuracies. However, there were some exceptional cases in which it was either infeasible or undesirable to record all the reported classification accuracies. For example, analyses that attempt to classify stimulus information at each TR over a window of time would be likely to yield highly correlated classification accuracies, due to the autocorrelation in fMRI BOLD signal. Therefore, in such cases, we only recorded the maximum decoding accuracy from the entire window. Another case concerns analyses conducted in both a single region and its constituent sub-regions, such as right, left, and bilateral dorsolateral PFC. We recorded only the sub-regions to reduce redundancy in our dataset. Finally, in cases where the goal of an analysis was to test whether the number of included voxels influenced the result, we again only recorded the maximum accuracy achieved. Thus, in general, we sought to include independent classification attempts in prefrontal cortex, and where classifications were non-independent, to favor inclusion of the one with the highest classification accuracy. Note that this latter inclusion criterion biases against the hypothesis that it is difficult to classify prefrontal cortex BOLD.

Another concern regarding the independence of observations relates to the intrinsic spatial smoothness of fMRI datasets. If decoding accuracies are obtained from two regions that are close enough to each other to show spatial correlation, the observations would not be independent. Given that papers do not report intrinsic spatial smoothness, we employed a threshold of 10mm as the minimum separation required for decoding accuracies to be considered separate. We found no cases of two analyses from the same study, decoding the same information, focused on regions that were less that 10mm apart. In addition, we conducted a second analysis where we averaged all accuracies from a study if they were associated with the same AAL region. This did not affect our results.

To ensure that classification accuracy values were reliably recorded from each paper we conducted a validation procedure in which an independent investigator (blinded to the initially coded value and without authorship incentive) re-coded the accuracy values from each paper. In each case where the two values were different, the values were re-checked and corrected in the record.

### Estimating Distributions

To compare decoding accuracy across regions (frontal v/s occipital or mid-temporal) and between significant and non-significant data we estimated distributions for the group mean data. First, the accuracies were pooled across studies and kernel density estimates were applied (using Scott’s rule for bandwidth). To obtain confidence intervals, we applied hierarchical bootstrapping which accounts for the dependence among analyses from the same study. Analyses from the same study share features that may influence classification accuracy such as sample size, data quality, preprocessing methods, etc. Studies were first randomly sampled with replacement and then group-level analyses were randomly sampled with replacement from these selected studies. Similar credible intervals were obtained by fitting Bayesian hierarchical Gaussian models to the data, though these models had to additionally assume a parametric family for the data.

In Study 70, mean decoding accuracies were reported in a bar graph and it was unclear whether decoding accuracies not significantly different from chance were excluded. Therefore, we excluded data from Study 70 from the estimation of the significant and non-significant distributions. Study 31 used 99.99% confidence intervals, reporting two accuracies at 63% and 64% within this interval. To be conservative, we did not count these accuracies in the estimation of the non-significant distribution.

### Regression Analysis

A regression analysis was employed to examine how decoding accuracy across the studies in our database depended on brain region within frontal cortex, the type of information decoded, and the analysis methods used. The coding of these factors for the regression is described below.

#### Region

We examined classification performance as a function of brain region. Individual analyses reported location using a number of different atlases. Therefore, to compare accuracy in a single space, we mapped all reported locations to the AAL atlas. Analyses that were reported with centroid coordinates, from either an ROI or a roaming searchlight were assigned to AAL regions by coordinate-lookup in SPM12’s AAL template image. Analyses that did not report coordinates used ROIs coming from one of several common brain atlases: Brodmann ^79^, Destrieux ^80^, Desikan-Kellaney ^81^, Oxford-Harvard (FSL). These analyses were assigned within AAL by a region-to-region correspondence table constructed by visually comparing the non-AAL atlases to the AAL atlas in MRICRON (Supplemental Table 2)

Within this region coding scheme, analyses with small ROIs or those that reported peak coordinates were each assigned to one AAL region, whereas analyses with larger ROIs were assigned to two or more AAL regions. 53%, 32%, 15% of the analyses were assigned to one, two, and three or more regions respectively. In most cases where accuracies were assigned to two AAL regions, these were the left and right hemisphere counterparts of the same AAL region. Therefore, for the main regression, analyses that had been assigned to two AAL regions were assigned to a bilateral region regressor. The accuracies not assigned to a bilateral ROI or assigned to more than 2 regions were omitted from the main analysis to maintain mutual exclusivity, leaving 82% of the original data. These accuracies were included in follow-up analyses described in the results and in the supplements.

#### Information Type

The type of information decoded in each analysis can be broadly categorized as either *perceptual, response, rule, or value.* We categorized an analysis as *perceptual* if trials were separated into classes so that they shared either a low-level perceptual feature such as color, or a high-level feature such as object category. Importantly, trials or patterns from the same class would be associated with different actions. In contrast, we categorized an analysis as *response* if trials from the same class contained the same action but different perceptual features. We categorized an analysis as *rule* if trials from the same class shared the same abstract goal, task or set of stimulus-response mappings. For example, one class of trials might require objects to be judged on their size, while the other class of trials might require judgments of shape. We considered an analysis as *value* if different classes of trials were associated with different levels of subjective value. For example, classes of trials might be distinguished based on a participant’s desire to purchase an object, or whether they experienced a win or loss outcome. Examples of each type of analysis can be seen in Supplementary Table 3.

#### Analysis Procedure

MVPA analyses varied along several dimensions at each step in the analysis pipeline from data collection and preprocessing to classification that could ultimately affect outcome. We recorded the following for each analysis at each step of the procedure: scanner strength, number of subjects, coregistration, smoothing, temporal averaging, response normalization, and classifier used. Specific codes used and their definitions are elaborated below.

##### Coregistration

Coded as 0-1 and refers to whether the decoding analysis was conducted in native-subject space or a standard space such as MNI.

##### Smoothing

Coded as 0-1 and refers to whether or not any smoothing kernel was applied to the fMRI data prior to the decoding analysis.

##### Temporal Averaging

Coded as one of four types, referring to how multiple fMRI images are combined into a single pattern corresponding to an experimental event. The four levels were:

1. *no temporal averaging*: uses every TR (repetition time) on every trial as a pattern
2. *averaging across trials*: averages data across trials, but maintains a separate pattern for each TR;
3. *averaging across TRs*: averages data across TRs but maintains a separate pattern for trial (or event)
4. *averaging across TRs and trials*: averages data across both trials and across TRs. This last category has the largest degree of temporal compression, often leading to only a few training examples per class.

##### Response normalization

Coded into 3 levels: no normalization, temporal normalization, and spatial normalization. Temporal normalization de-means each voxel across time and divides by the standard deviation either within class or across all classes. Examples using this method are Studies 36, 25, 4. Spatial normalization de-means each voxel using the average response of the surrounding voxels. An example using this method is Study 3.

##### Classifiers

Coded as one of 6 types: Gaussian naive Bayes (gnb), logistic regression (logreg), linear discriminant analysis (lda), linear support vector machines (svm-lin), nonlinear support vector machines (svm-nonlin), and correlation (correlation). Correlation accuracy is determined by assessing whether the within class correlation is higher than the between class correlation in random splits of the data.

### In-house fMRI dataset

In order to complement our meta-analysis, we also analyzed previously collected fMRI data from our laboratory on a cognitive control task requiring coding of rule and visual stimulus information. Rule information was indicated by a visual feature, so both types of information could be decoded from the same set of trials. This allowed us to compare decoding accuracy between frontal and visual cortex, while controlling for differences in methods. We also employed this dataset to compare the signal-to-noise ratios and pattern reliability between frontal and visual cortex within this dataset. The details of the task and analyses can be found in the Supplementary Materials.

## Author Contributions

A.B., C.G. & D.B. designed the study. C.G. & A.B. carried out the analyses. A.B., C.G. & D.B. wrote the paper. A.B. & C.G. contributed equally to this work.

## Acknowledgements

We are grateful to Marc Coutanche & colleagues for kindly sharing their meta-analysis data of visual decoding studies. We thank Brittany Ciullo, Nada Hamzah, Juliana Trach, Sarah Master and Celia Ford for their assistance with independently verifying the coding of the meta-analysis data. This work was support by grants from NINDS (NS065046) and NIMH (MH099078, MH111737) at the NIH, and a MURI award from the Office of Naval Research (N00014-16-1-2832).

